# Multisensory integration while learning to read: A longitudinal functional and structural MRI study

**DOI:** 10.64898/2026.07.24.740626

**Authors:** Johanna Finnemann, Ga-ram Jeong, Tzipi Horowitz-Kraus, Michael A. Skeide

## Abstract

Literacy development involves the recognition and integration of auditory and visual signals and as such offers a unique opportunity to track the development of an eventually efficient multisensory processing network.

To shed light on this topic we conducted a 12-month longitudinal 3 Tesla functional and structural MRI study in children beginning to receive formal literacy instruction. Twenty-three six-year-old German-speaking participants completed four fMRI sessions during which they were presented with letters, speech sounds, or their audiovisual combinations in congruent and incongruent formats. Longitudinal changes in BOLD response amplitude were assessed using linear mixed-effects models. Over the course of learning to read, we observed significant changes in modulations of BOLD activity for the ‘congruent - incongruent’ contrast within a left-lateralised network encompassing the central opercular cortex, anterior supramarginal gyrus, inferior frontal gyrus (pars triangularis) and frontal pole. These effects were primarily driven by a gradual increase of response amplitude across sessions, consistent with the refinement of audiovisual integration mechanisms. Structural analyses revealed longitudinal anatomical variation within temporal, occipital, and sensorimotor regions, predominantly involving the right middle and inferior temporal gyri and bilateral temporal regions. However, changes in regional anatomy and functional activation were not significantly associated with behavioural measures of grapheme-phoneme association. Together, these findings demonstrate reorganisation within established regions known to be related to language processing and multisensory integration during early literacy development, highlighting the utility of longitudinal MRI for tracking experience-dependent neuroplastic changes in brain function and structure.

## Introduction

Perception begins with a riot of sensory signals: fragmented sights, sounds, textures, and smells that bear no intrinsic markers of their origin. As James famously described, the world initially appears as a “blooming, buzzing confusion”, and the task of the developing brain is to carve structure from this influx. Because objects in the environment typically generate multiple, simultaneous sensory cues, a central challenge of perception is to infer which signals arise from the same underlying cause. The coherence of even the simplest object such as the form and colour of a pebble as Bain noted back in 1855, the feel of its surface, and the sound it makes when struck, depends on binding disparate sensory impressions into a unified representation (Bain 1855). Mature perceptual systems are fairly efficient at solving most of these problems, integrating information across modalities to construct stable, causally organised models of the world.

From a computational perspective, multisensory perception is therefore an exercise in causal inference: determining whether co-occurring cues should be combined or treated as independent. Developmental theories have long debated how such integrative abilities arise. Early theoretical accounts have contrasted differentiation frameworks, which propose that infants begin with broad, undifferentiated multisensory representations that later fractionate into modality-specific systems, with integration frameworks, which posit that efficient multisensory binding emerges only after sufficient unisensory maturation (Burr & Gori, 2012). While infants show early sensitivity to rudimentary cross-modal correspondences i.e. spatial and temporal congruency detection (Lewkowicz & Kraebel, 2004), more sophisticated forms of cue combination such as weighting inputs by their relative reliability or incorporating prior knowledge (Gau & Noppeney, 2016; Tong et al., 2020) about environmental regularities emerge gradually over childhood. This protracted trajectory suggests that multisensory integration is refined through a combination of neural maturation, experience-dependent calibration, and the increasing availability of top-down expectations based on statistical regularities (Gori et al., 2008). Consequently, childhood represents a period of heightened plasticity in which multisensory computations transition from being predominantly stimulus-driven to increasingly flexible, and context-sensitive.

These developmental dynamics are especially salient in the domain of literacy. Learning grapheme-phoneme correspondences requires both the creation and the continual refinement of bidirectional audiovisual associations, enabling readers to activate phonological codes from print and, conversely, to evoke orthographic information from speech (Regev et al., 2013). Crucially, this reciprocity is not supported by any natural co-occurrence structure: letters and phonemes do not arise from a common physical source, and the brain must impose this linkage through explicit instruction and cumulative experience. With practice over an extended period of time, these associations become automated, and the integration of letters and sounds begins to exhibit the efficiency and immediacy characteristic of naturalistic multisensory processing (Raij et al. 2000; Blau et al., 2008; Blomert & Froyen, 2010). Tracing how the neural architecture supporting this integration reorganises over the course of reading acquisition provides a window into experience-driven plasticity in the developing brain.

The neural circuitry supporting these multisensory and language-related computations spans a broad left-lateralised network that continues to reorganise throughout early childhood. Regions along the superior temporal and temporo-parietal cortices contribute to mapping speech sounds onto their visual or articulatory counterparts, whereas inferior parietal and inferior frontal areas support phonological manipulation, conflict resolution, and the evaluation of audiovisual congruency (Beauchamp et al., 2010; Morís Fernández et al., 2017; Goranskaya et al., 2016; Kita et al., 2013; Kolozsvári et al., 2019). Frontal regions, including mediofrontal territories, are implicated in exerting top-down control over newly learned associations and in monitoring mismatches between expected and perceived cues (Garrison et al., 2013; Schlichting & Preston, 2015). Together, these areas form a distributed system that gradually becomes more efficient, more specialised, and more automatic as children become fluent readers.

From an imaging perspective multisensory integration has traditionally been operationalised using superadditivity, i.e. the idea that responses to multisensory stimuli should exceed the sum of responses to their unisensory components (Calvert et al., 2000; Calvert et al., 2001; Stevenson et al., 2007; Wright et al., 2003), however this criterion is increasingly recognised as overly conservative for BOLD imaging because vascular nonlinearities and mixed neuronal populations can obscure integrative responses (Beauchamp, 2005; Laurienti et al., 2005; Goebel & van Atteveldt, 2009). Consequently, studies of learned audiovisual associations such as letter–speech sound integration commonly assess the neural discrimination of congruent and incongruent audiovisual pairings, an approach that avoids many of the limitations associated with unisensory subtraction (Goebel & Van Atteveldt, 2009). We therefore focused primarily on congruency-related responses as a measure of audiovisual letter–speech sound integration, consistent with previous fMRI studies that have used audiovisual congruency to probe multisensory integration (Blau et al., 2008; Hein et al., 2007; Noppeney et al., 2008; van Atteveldt et al., 2004; van Atteveldt et al., 2007), while additionally examining superadditive responses as a complementary measure.

Although cross-sectional work has identified components of this network in children at varying stages of reading proficiency, little is known about how these multisensory integration and language regions change over time within the same individuals, or how they acquire the ability to discriminate congruent from incongruent letter–sound pairings as reading skills mature.

Longitudinal studies are particularly well suited to addressing these questions, as they allow the unfolding of multisensory and language-related specialisation to be tracked across the period in which children transition from prereaders to emerging readers. Yet relatively few MRI investigations have captured neural development over the first year of formal literacy instruction, a window during which behavioural measures show rapid gains in explicit letter–sound knowledge and phonological awareness. A comprehensive longitudinal characterisation of both regional response strength and distributed activation patterns is therefore essential for understanding how the brain’s multisensory– language system becomes tuned to the demands of print–speech integration at the early stages of learning to read.

## Results

### Behavioural results

In a behavioural task performed offline outside of the scanner children completed an audiovisual congruency task in which they judged whether visually presented letters and simultaneously presented speech sounds were congruent or incongruent. Full task details are provided in the Methods. Task performance, in particular drift rate (*v*) increased significantly across the four sessions (β = 0.375, SE = 0.065, *z* = 5.74, *p* < .001), indicating progressively more efficient evidence accumulation over the first year of literacy instruction. The estimated intercept was 0.664 (SE = 0.193, *p* < .001) (Figure 1). In contrast, boundary separation (a) did not change significantly across sessions (β = 0.025, SE = 0.056, z = 0.442, *p* = 0.636), suggesting that participants maintained a stable response criterion throughout the study. The estimated intercept was 1.220 (SE = 0.152, *p* < 0.001).

**Figure 1.**
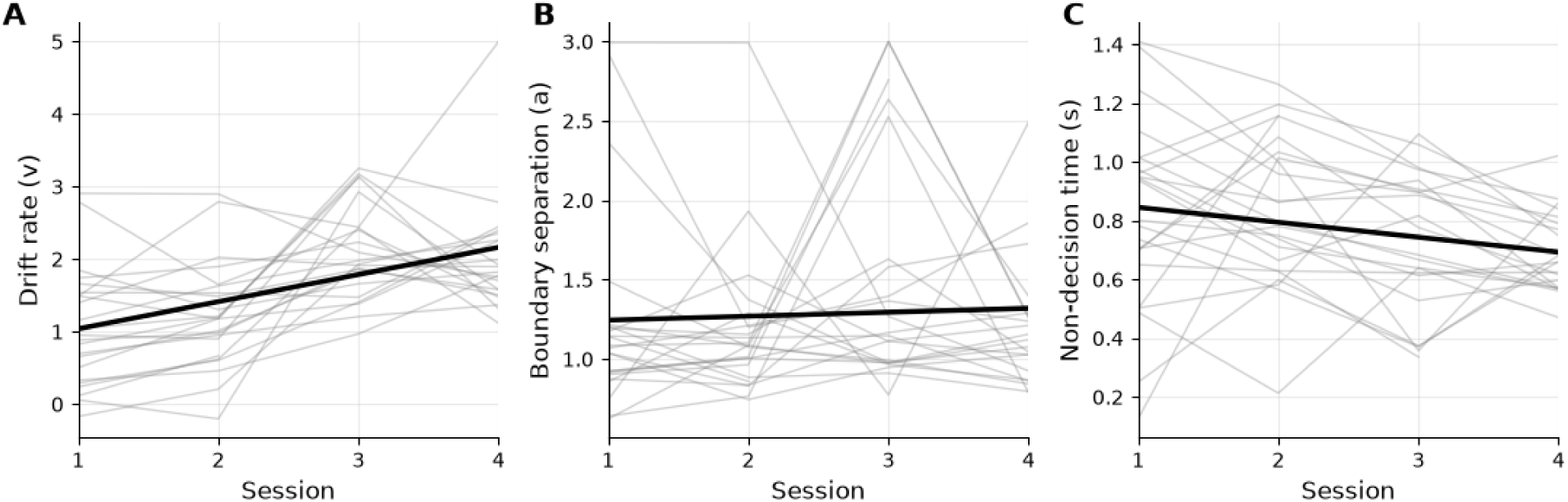
Session-related changes in drift diffusion model parameters. Drift diffusion model (DDM) parameters were examined across four experimental sessions using linear mixed-effects models with session as a fixed effect and participant as a random intercept. Individual participant trajectories are shown in grey. (A) Drift rate (*v*) reflects the efficiency of evidence accumulation during decision-making. (B) Boundary separation (*a*) reflects response caution and the amount of evidence required before committing to a decision. (C) Non-decision time (*s*) reflects processes unrelated to evidence accumulation, including perceptual encoding and motor response execution.

Non-decision time (s) decreased significantly across sessions (β = −0.051, SE = 0.020, z = −2.55, *p* = 0.011), indicating faster non-decisional processes, such as stimulus encoding and motor execution, over time. The estimated intercept was 0.896 (SE = 0.0561, *p* < 0.001).

Taken together, these findings suggest that longitudinal improvements in task performance were driven primarily by more efficient evidence accumulation and faster non-decision processes related to the maturation of sensory and motor processing, rather than changes in response caution.

### Functional imaging results

To identify how brain areas processing orthographic and phonological alignment reorganise when learning to read, we evaluated the longitudinal fixed effect of time on the ‘congruent - incongruent’ contrast using mass-univariate linear mixed-effects models (LMMs).

Significant increases over time for the ‘*congruent - incongruent*’ contrast emerged across left frontoparietal and perisylvian areas with peak clusters located in the anterior supramarginal gyrus, frontal pole, central opercular cortex and inferior frontal gyrus (pars triangularis) (Figure 2 and Table 1). The longitudinal changes in activation within the functional ROIs were not significantly associated with changes in behaviour (drift rate - *v*) over time. For a detailed overview of these results see S1. To complement the primary congruency analyses, we also examined the classical superadditivity contrast (*congruent-minus- (unimodal auditory + unimodal visual)*), which has traditionally been used as a stringent fMRI measure of multisensory integration. No significant longitudinal changes were observed (all *p* > 0.05 FEW-corrected).

**Figure 2.**
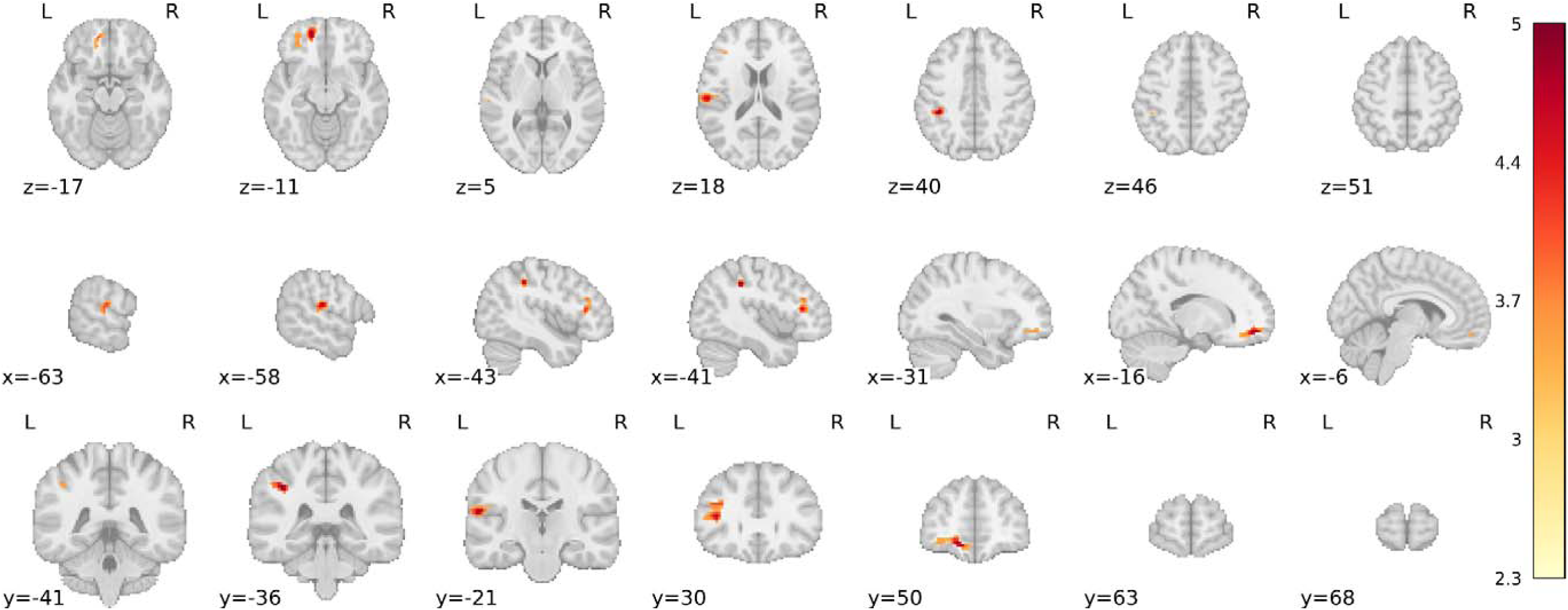
Longitudinal group-level linear mixed-effects analysis of the congruent-minus-incongruent contrast showing time-sensitive increases in the neural congruency effect during literacy acquisition. Statistical maps are shown in MNI152NLin6Asym standard space and corrected for multiple comparisons using two-sided cluster correction (voxel-wise threshold *p* < 0.001; cluster-level *p* < 0.05). Maps are displayed at |Z| ≥ 2.3 with clusters ≥14 voxels.

**Table 1.**
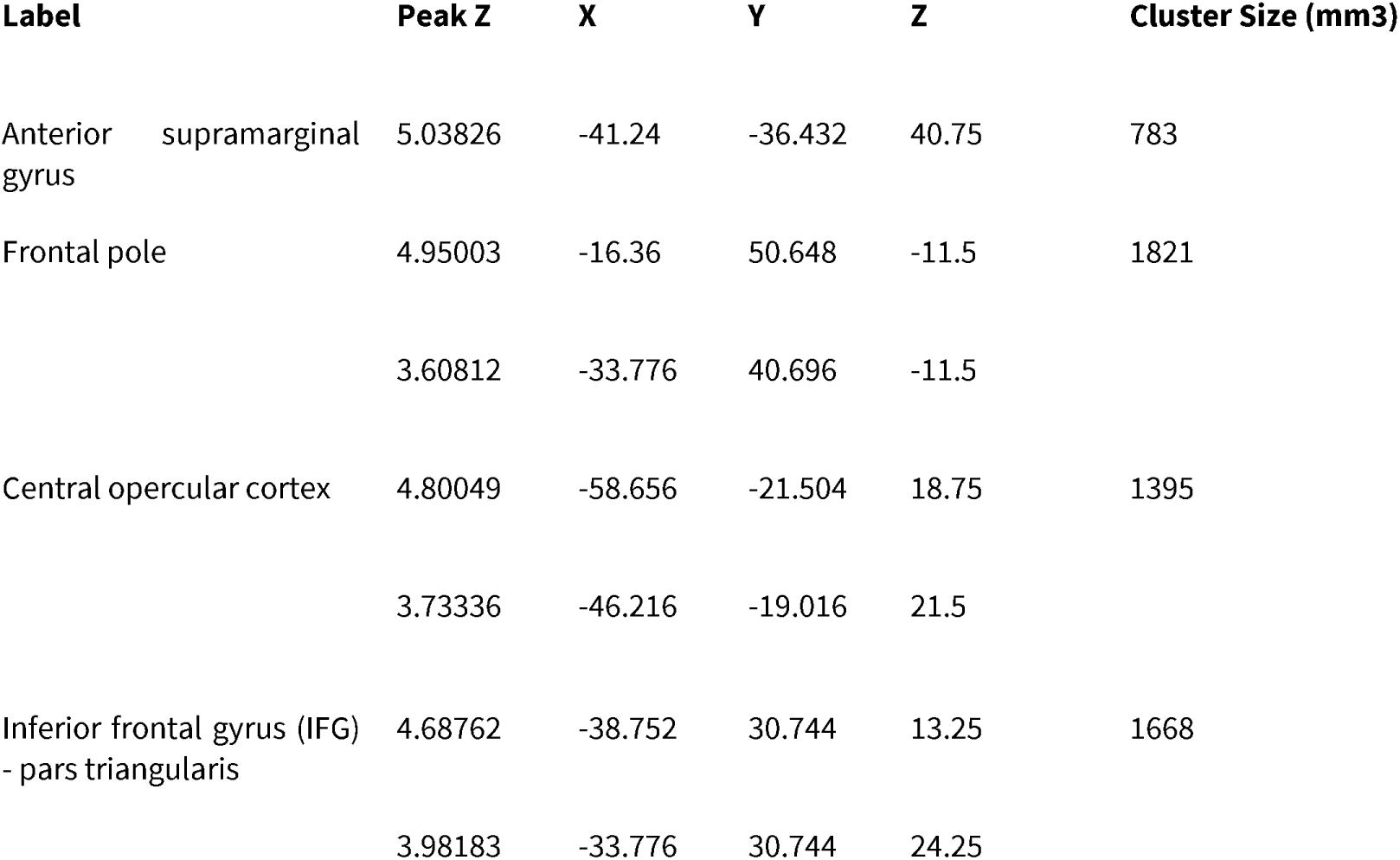
Significant activation clusters for the congruent-minus-incongruent contrast. Coordinates are in standard MNI space and anatomical labels were assigned using the Harvard–Oxford probabilistic atlas (implemented in *FSL/Nilearn*).

| Label | Peak Z | X | Y | Z | Cluster Size (mm3) |
| --- | --- | --- | --- | --- | --- |
| Anterior supramarginal gyrus | 5.03826 | -41.24 | -36.432 | 40.75 | 783 |
| Frontal pole | 4.95003 | -16.36 | 50.648 | -11.5 | 1821 |
|  | 3.60812 | -33.776 | 40.696 | -11.5 |  |
| Central opercular cortex | 4.80049 | -58.656 | -21.504 | 18.75 | 1395 |
|  | 3.73336 | -46.216 | -19.016 | 21.5 |  |
| Inferior frontal gyrus (IFG) - pars triangularis | 4.68762 | -38.752 | 30.744 | 13.25 | 1668 |
|  | 3.98183 | -33.776 | 30.744 | 24.25 |  |

For percent signal change plots over the four sessions for each significant cluster and single contrast plots see S2.

### Structural Imaging

In addition to functional changes, longitudinal MRI enables the characterisation of structural maturation over the same developmental period. Measures of surface area, curvature and cortical volume undergo substantial age-related changes throughout childhood and may provide complementary information about the anatomical maturation of regions supporting emerging literacy.

The vertex-wise linear mixed effects analysis revealed distinct patterns of structural surface changes across the tracking period, see Table 2 and Figure 3. These longitudinal changes in cortical morphology were not significantly associated with drift rate in either the linear mixed-effects (LME) or ordinary least-squares (OLS) linear regression with the exception of the sixth curvature cluster where baseline curvature positively predicted increase in drift rate (β = 0.523, 95% CI [0.194, 0.852], p = 0.0399, FDR-corrected). For a summary of the LME and OLS results see S3 and S4. Coordinate-based functional decoding of the relevant curvature cluster using Neurosynth (Yarkoni et al., 2011) indicated associations with studies involving developmental studies of phonological processing and reading (McNorgan et al., 2014; Meyler et al., 2008), articulatory accuracy of phonemes associated with cortical thickness (Tremblay & Deschamps, 2016), phonological judgment (Zhao et al., 2014) as well as conflict resolution during congruency judgment tasks (Floden et al., 2012; Schulte et al., 2011).

**Figure 3.**
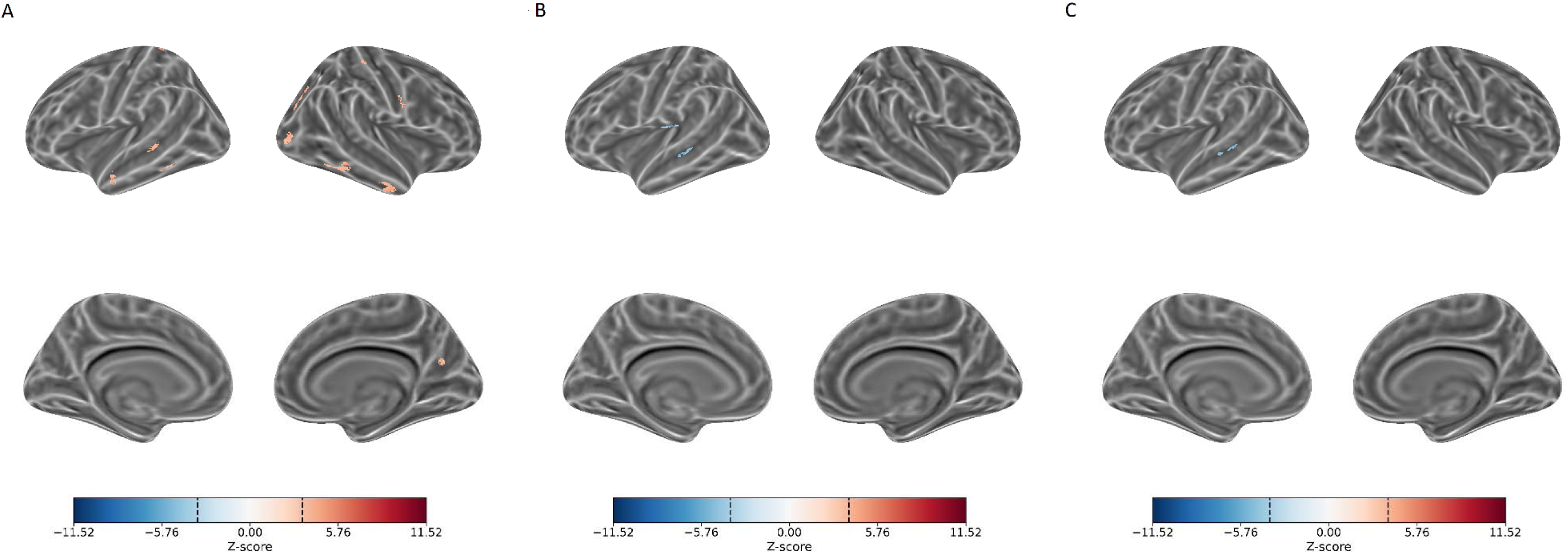
Longitudinal changes in cortical surface morphology. Changes in curvature, surface area and grey matter volume are projected onto the inflated *fsaverage* template across lateral, medial, dorsal, and ventral orientations for both the left and right hemispheres for (A) Cortical Curvature, (B) Cortical Surface Area and (C) Cortical Volume. For each metric, the z-score threshold was determined using the calculated p-value corresponding to a false discovery rate threshold of *p_FDR_* < 0.05, corresponding to a threshold of ±3.420 for curvature, ±3.875 for cortical surface area and ±3.869 for volume. The cluster-extent threshold was surface area > 50mm^2^.

**Table 2.**
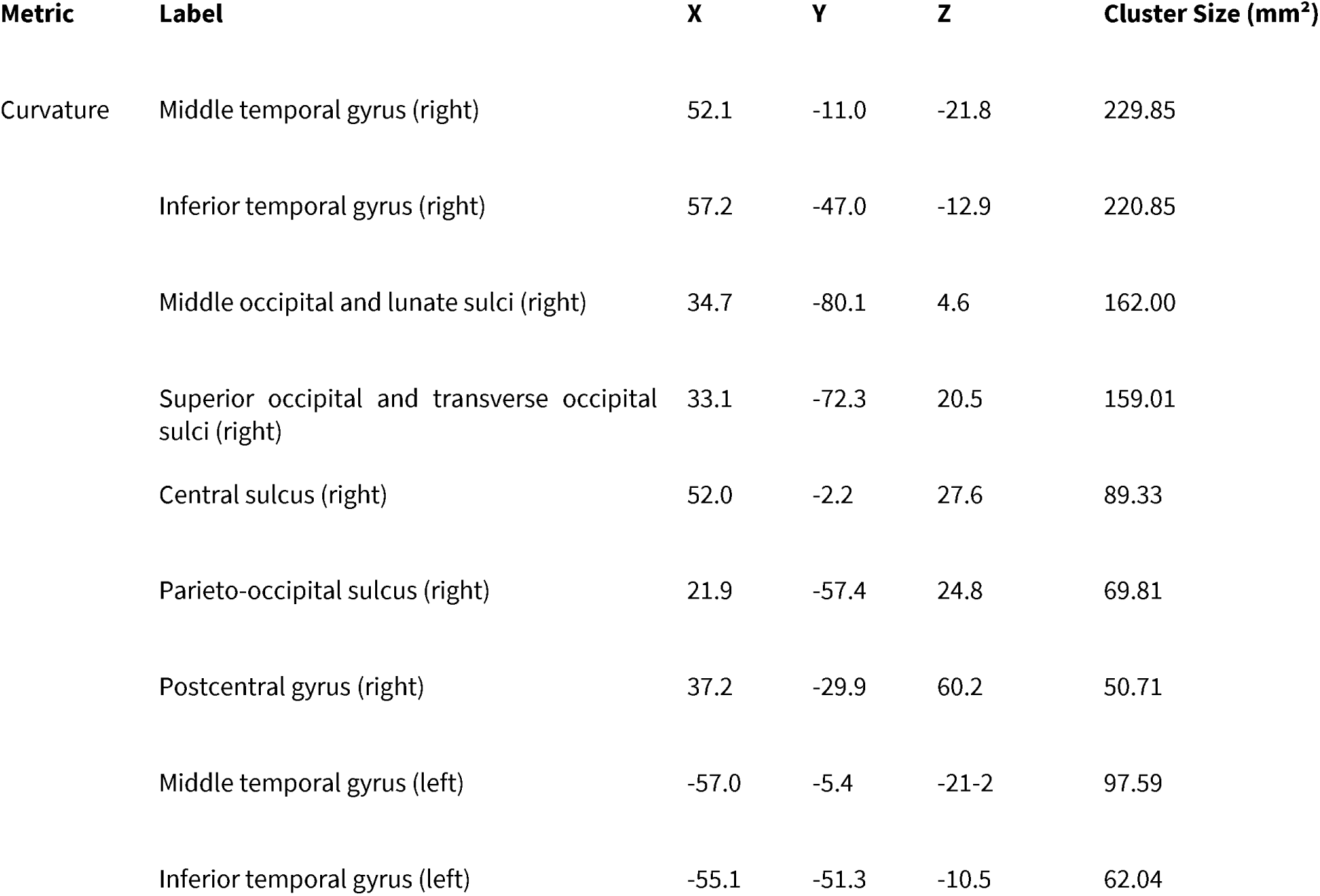
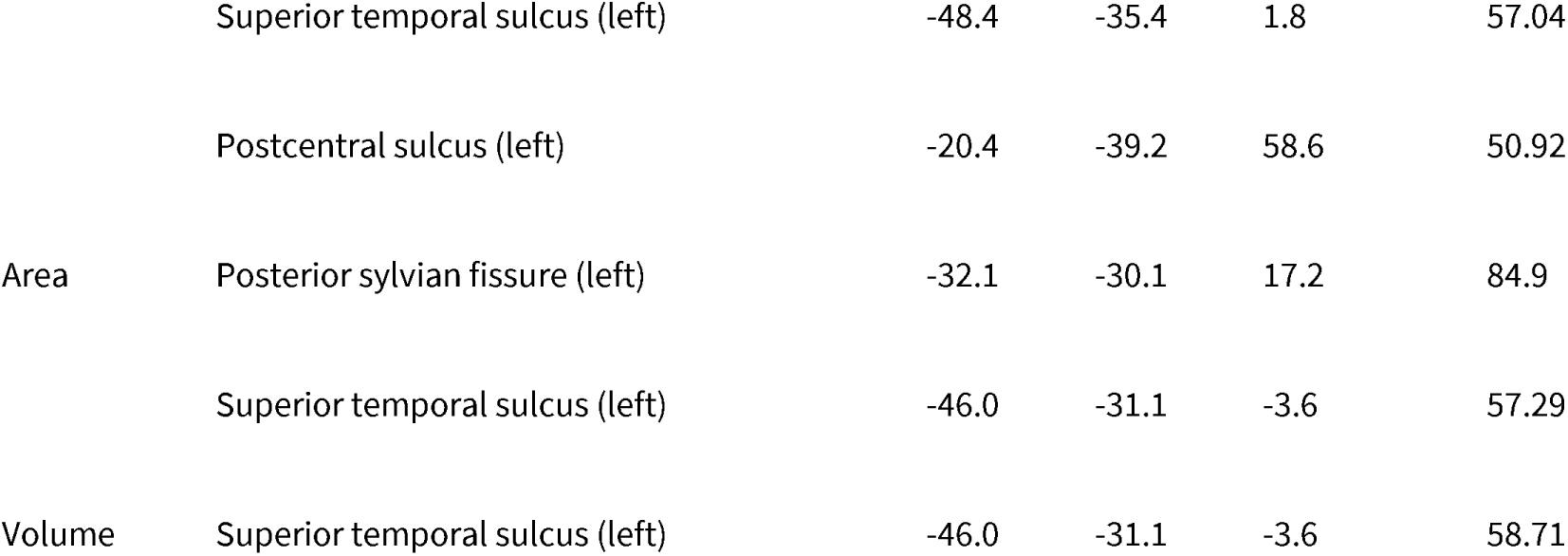
Significant longitudinal changes in cortical surface morphology. Coordinates are in MNI 305 space and anatomical labels were assigned using the Destrieux atlas (a2009s).

To assess the spatial correspondence between functional and structural changes, we overlaid the functionally derived cluster masks from the second-level analysis onto the structural results and examined whether any significant structural effects were located within these regions. No overlapping significant voxels were identified.

## Discussion

We investigated how behavioural performance, functional responses to letter–speech sound integration, and cortical morphology change during the first year of formal reading instruction. Behaviourally, children showed increasingly efficient letter–speech sound processing and functionally, congruency-related responses increased over time within a distributed left-lateralised fronto-parietal network implicated in audiovisual and phonological processing. No longitudinal changes were observed for the classical superadditivity contrast. In addition, several cortical regions exhibited longitudinal morphological changes consistent with ongoing structural maturation with higher baseline curvature in the parieto-occipital cortex/precuneus predicting greater literacy gains on the behavioural task.

The longitudinal improvements in the 2-alternative-forced-choice letter–phoneme congruency judgment task were characterised by a significant increase in drift rate across sessions, indicating progressively more efficient evidence accumulation for congruent versus incongruent mappings. This finding suggests that children became increasingly effective at extracting and integrating relevant audiovisual information over the first year of reading instruction. In contrast, boundary separation remained stable over time, indicating that response caution or decision thresholding did not systematically change across sessions. This pattern implies that improved performance was not driven by strategic shifts in speed–accuracy trade-offs, but rather by changes in information processing efficiency.

In addition, non-decision time decreased significantly across sessions, suggesting faster peripheral processing components, including improved stimulus encoding and more efficient motor execution. Together, these effects indicate that longitudinal gains were primarily driven by enhancements in both evidence accumulation and basic processing speed, consistent with developmental refinement of task-relevant perceptual and sensorimotor processes rather than changes in decision strategy.

Across the first year of formal reading instruction, increasing activation for congruent relative to incongruent letter–sound pairings was observed in a distributed left-lateralised network encompassing the parietal operculum, anterior supramarginal gyrus, inferior frontal gyrus (IFG), and frontal pole. Activity increases in the parietal operculum and anterior supramarginal gyrus likely reflect strengthening auditory–orthographic binding mechanisms and improved phonological decoding efficiency. The engagement of inferior frontal regions and the frontal pole suggests increasing reliance on top-down language control during cross-modal integration. These changes may index enhanced predictive or selection mechanisms that support the resolution of competing phonological representations during reading, rather than basic sensory convergence, especially when decisions about possible congruency have to be made. Increased activation in the left pars triangularis may be associated with the growing recruitment of phonological and other language-related processes during grapheme-phoneme matching, potentially supporting the retrieval and evaluation of learned letter-sound associations.

Taken together, these findings suggest that learning to read strengthens a left-lateralised fronto-parietal network underlying controlled language processing (Qi et al., 2021; Skeide & Friederici, 2016). Alongside these functional changes, longitudinal morphological changes were observed across a broader set of cortical regions including bilateral temporal, occipital, parieto-occipital and sensorimotor areas. On the whole there was no relationship between changes in cortical morphology and behavioural improvement with the exception of baseline curvature in the cluster encompassing the parieto-occipital sulcus/precuneus where higher curvature predicted subsequent increases in drift rate. This raises the possibility that greater baseline cortical maturity in this region may support more efficient phonological processing and conflict resolution thereby facilitating the acquisition of phoneme-grapheme mappings during early reading development.

A key constraint for the interpretation of the current findings is that grapheme–phoneme integration may not be fully equivalent to other forms of audiovisual multisensory integration (e.g., audiovisual speech or temporally congruent arbitrary pairings). As highlighted by Blomert & Froyen (2010), literacy-related mappings may recruit partially distinct neurobiological mechanisms, potentially due to their symbolic, culturally learned nature and explicit instruction. Consequently, the observed activation changes may reflect domain-specific tuning of learned orthographic–phonological associations rather than canonical multisensory integration processes.

In addition, children were not completely naïve to letters and their associated speech sounds at the beginning of the study and entered formal instruction with varying levels of pre-existing alphabetic knowledge. A related limitation is the absence of a control group that did not undergo formal reading instruction over the same period. As the first year of schooling coincides with substantial changes in cognitive and attentional demands, as well as cortical development any longitudinal changes in behavioural performance and functional responses may partially reflect more general developmental processes. Thus, although the observed changes in congruency-related activity are consistent with increased efficacy of grapheme-phoneme associations the present study does not allow these effects to be uniquely attributed to reading instruction over the impact of task familiarity or general cognitive development.

Finally, the reported developmental increases may also reflect progressive refinement of attentional or layer processing streams, consistent with proposals that sensory integration and top-down modulation can be partially dissociable at microcircuit levels (e.g., Gau et al., 2020), with early literacy learning progressively shifting processing possibly toward more efficient, feedback-supported integration (Skeide & Friederici, 2016). These distinct depth-dependent profiles suggest that multisensory and attentional mechanisms may regulate sensory processing via partly distinct circuitries. However, in the present study these processes cannot be disentangled, as the 3 Tesla fMRI data are not sensitive to cortical layer-specific activity, and thus do not allow inferences about laminar-level activity.

## Methods

### Participants

A total of 32 participants were recruited consisting of children who entered primary school in the summer of 2023. Telephone screening was conducted with each family: German was required to be their first and only language, in addition they should not have any hearing or visual impairment(s), no developmental disorders and not take any regular medication. Parents were asked about their own education, household income and their children’s academic abilities. Compared to 100 children of the same age, parents ranked their children on average as coming 43^rd^ and 41^st^ in writing and reading abilities respectively.

All children took part in an initial mockup session, where they were introduced to a mock scanner, could lie down to see what it feels like and were played scanner sounds. If they were comfortable with the procedure they were invited back for their first scan. To increase motivation each child was provided with a little ‘research booklet’ in which they could earn stickers for each attendance and upon completion of the study received a small present from the study team. One child did not return for the first MRI session and a further 5 decided not to continue with the study after the first MRI session yielding 26 participants who completed 2 or more sessions.

Following MRI preprocessing and quality control, three additional datasets were excluded, resulting in a final sample of 23 children included in the analyses. The final analytic sample had a mean age of 6.46 years at intake (SD = 0.24), with an equal gender distribution (12 females, 11 males). The mean interval between MRI sessions in the final sample was 113.37 days (SD = 45.95).

All methods were carried out in accordance with the Declaration of Helsinki and all experimental protocols were approved by the Ethics Committee of the Medical Faculty of the University of Leipzig, Germany (438/20-ek). Informed consent was obtained from a parent, and the children gave documented verbal assent to participate in the study.

### Behavioural Task: Stimulus and Task Design

Outside of the scanner participants first completed a practice session designed to familiarise them with the task design. The experiment that was based on the design introduced by van Atteveldt, et al. 2004 was programmed and executed using *PsychoPy* (version 2022.2.5). In each mini-block, a sequence of six images was presented while corresponding sounds were played simultaneously. Trials were either congruent, where the visual and auditory stimuli matched, or incongruent, where the auditory stimulus did not correspond to the displayed image. Participants judged whether the audiovisual pairing was correct (congruent) or incorrect (incongruent) using designated response keys (‘K’ for correct and ‘D’ for incorrect). Immediate feedback was provided after each response to facilitate familiarisation with the task. In the practice experiment the images were six cartoon animals (cat, chicken, dog, goat, horse, lion) and their respective sounds (meow, cluck, bark, bleat, neigh, roar). The practice run included 24 mini-blocks pseudorandomised and counterbalanced across congruent and incongruent pairings, followed by 48 grapheme-sound mini-blocks: Stimuli were speech sounds corresponding to single consonants and their visually presented upper-case single letters: B, F, K, M, P and T again counterbalanced and pseudorandomised.

Each mini-block consisted of a rapid, rhythmic stream consisting of six successive presentations of the same stimulus pairing with the audio-visual stimulus being presented for 467ms followed by an inter-stimulus interval of 200ms. The maximum length of each mini-block was 6 seconds including a ∼2 second response window at the end. Both accuracy and response times were recorded.

Results were analysed using drift diffusion modelling (DDM) implemented in *Python* (version 3.12.9) using the *PyDDM* package. Reaction times (RTs) shorter than 200ms or longer than 5s were excluded prior to analysis. DDM parameters were estimated separately for each participant and session by fitting a standard drift diffusion model to the joint distribution of response accuracy and reaction times using maximum-likelihood estimation. The model included three free parameters: drift rate (*v*), boundary separation (*a*), and non-decision time (*s*). Parameter search ranges were specified as follows: drift rate (*v*) = −5 to 5, boundary separation (*a*) = 0.3 to 3, and non-decision time (*s*) = 0.1 to 3 s. To assess longitudinal changes in decision-making processes, linear mixed-effects models were fitted separately for drift rate, boundary separation, and non-decision time using session number as a fixed effect and participant as a random intercept.

Inside the scanner 12 grapheme-sound mini-blocks were presented per run in the same way as before yielding a total of 48 mini-blocks across 4 runs, but participants were only instructed to attend to the pairings without responding to them to prevent strong (pre)motor cortex activation and motion artifacts. As a basic attention check during the scanning session children were instructed to press a button whenever they heard the sound of a bell (instead of the phonemes) or saw a picture (instead of the visual letters). Accuracy for the attention check was 90.8% across all subjects and sessions indicating overall very good attention.

### Imaging Data Acquisition

MRI data were acquired on a Siemens MAGNETOM Prisma 3T scanner at the Max Planck Institute for Human Cognitive and Brain Sciences using a 64-channel head coil. Following an initial localiser scan, participants completed four functional runs. If head motion was deemed excessive and the child remained compliant, an additional functional run was acquired. Functional images were collected using a multiband gradient-echo echo-planar imaging (EPI) sequence (TR = 2000ms, TE = 22ms, flip angle = 80°, field of view = 204 mm, voxel size = 2.5 × 2.5 × 2.5 mm, 60 axial slices, multiband factor = 2, anterior–posterior phase encoding). Each functional run consisted of 130 volumes.

At the end of the session, a high-resolution T1-weighted structural image was acquired using an MPRAGE sequence (TR = 2300ms, TE = 3.26ms, TI = 900ms, flip angle = 9°, field of view = 256 mm, 1 mm isotropic voxels, 176 slices). During this structural scan, children were allowed to watch a video of their choice to facilitate compliance. In the first wave of data collection, an additional FLAIR sequence was acquired and reviewed by study physicians to screen for incidental findings.

### Data preprocessing

All imaging data were stored and processed on a server located at the Max-Planck-Institute for Human Cognitive and Brain Sciences. Data were BIDS-converted and defaced using a locally installed version of *ezBIDS* (Levitas et al., 2024) with *pyDeface* (Gulban et al., 2019). *MRIQC* (Esteban et al., 2017) was run to generate standardised image quality metrics and visual reports for transparency. These outputs were inspected but not used as an additional decision layer beyond predefined quality control rules. Preprocessing was performed using *fMRIPrep 24.1.1* (Esteban et al., 2018 & 2020). Data were processed using the longitudinal flag which constructs a within-subject template to improve anatomical consistency across sessions and reduces bias in structural alignment.

Structural processing was performed using *FreeSurfer*, including longitudinal stream reconstruction for subjects with multiple sessions. This approach generates a subject-specific unbiased template and propagates anatomical information across timepoints to improve robustness of cortical reconstruction and registration.

### Data exclusion and Noise Regression

One participant was excluded from the analysis due to the presence of a temporal arachnoid cyst. Runs were excluded if more than 30% of volumes exceeded a framewise displacement threshold of 0.7 mm (Power et al., 2012). Outlier time points and non-steady-state volumes were identified and included as nuisance regressors in the first-level models. In addition, regression parameters for the functional volumes included the 6 canonical head motion parameters as well as the first 6 *CompCor* components to account for physiological noise (Behzadi et al., 2008).

Only sessions with successfully estimated first-level models were entered into group-level analyses. This yielded a total of 23 participants with 82 valid sessions and 297 runs. For a more detailed overview of the participants, sessions and runs please refer to supplementary Table S5.

Subject-level and session-level statistical analyses were implemented using a General Linear Model (GLM) framework via *Nilearn*. Prior to model estimation, functional images were slice time corrected, co-registered to the subject’s skull-stripped averaged anatomical image and warped to the MNI152NLin6Asym template. Functional volumes were spatially smoothed using a 7.0mm Full-Width at Half-Maximum (FWHM) Gaussian kernel.

The BOLD response was modelled using the canonical Glover Hemodynamic Response Function (HRF) combined with its temporal derivative and dispersion terms to capture variations in response onset and duration. A comprehensive set of directional contrasts was specified to extract beta weights for individual conditions and condition differences: *unimodal auditory, unimodal visual, congruent, incongruent, congruent-minus-unimodal auditory, congruent-minus-unimodal visual, congruent-minus-(unimodal auditory + unimodal visual)* and *congruent-minus-incongruent*. The *congruent-minus-incongruent* contrast was the main contrast of interest indexing sensitivity to learned grapheme– phoneme correspondences. The *congruent−minus-(unimodal auditory+unimodal visual)* contrast was additionally examined as a measure of superadditivity, defined as an enhanced audiovisual response that exceeds the combined response predicted from the two constituent unisensory inputs.

Group-level statistical inference was performed using voxel-wise Linear Mixed-Effects (LME) models implemented in *Julia*. A mixed model was fitted to the extracted first-level contrast effect size maps using the following formula:

Beta ∼ Time + (Time|Subject)

To control the family-wise error (FWE) rate while preserving statistical power, a cluster-extent correction threshold was derived via spatial autocorrelation function (ACF) modelling.

We used the parametric cluster correction suggested by Cox et al. (2017) and implemented in *AFNI*’s (version 24.3.08) *3dFWHMx* and *3dClustSim* programme which models non-Gaussian noise (Cox et al., 1996, 1997; Eklund et al. 2016). Statistical maps were thresholded at a voxel-wise cluster-defining threshold of *p* < 0.001 with a two-sided cluster-level family-wise error (FEW) threshold of *p* < 0.05. Using 10,000 Monte Carlo simulations, this corresponded to a minimum cluster extent of 28 voxels. Only clusters exceeding this were reported.

To examine whether neural changes tracked behavioural improvement across sessions, cluster-wise mean activation values were extracted from the significant clusters identified in the primary whole-brain analysis and related to the drift diffusion model drift rate (*v*). Separate linear mixed-effects models were fitted for each cluster with the session × *v* interaction as the effect of interest. P-values for the interaction term were corrected for multiple comparisons across clusters using false discovery rate (FDR).

### Structural MRI Analysis

We also analysed longitudinal changes in cortical structure using surface-based morphometric data derived from *FreeSurfer* reconstructions. Subject-level and session-level statistical analyses were implemented using the same formula as for the volumetric functional analyses.

T1-weighted structural MRI data were processed using the longitudinal stream in *FreeSurfer* (version 7.4.1). To extract reliable anatomical metrics across multiple timepoints and minimize confounding cross-sectional biases, a three-step longitudinal processing pipeline was implemented. First, every individual session (up to four sessions per participant) was independently processed through the standard recon-all pipeline for initial motion correction, skull-stripping, intensity normalisation, and automated topology tiling. Second, an unbiased, within-subject template was constructed for each participant using *recon-all -base*. This step combines anatomical data across all available timepoints to create a subject-specific space that acts as a common template. Each independent session was re-processed through the *recon-all -long* workflow, where information from the within-subject template was utilised to guide surface registration, refine tissue boundaries (e.g., grey/white matter borders), and dramatically reduce measurement noise over time. Finally, surface data were spatially smoothed using a Gaussian kernel with a full-width at half-maximum (FWHM) of 5mm for the area and volume-based metrics and 7mm for the curvature metric to optimise the signal-to-noise ratio. Vertex-wise statistical analyses were performed within a Linear Mixed Effects (LME) framework using *FreeSurfer*’s *MATLAB* toolkits. To comprehensively evaluate structural changes, separate LME models were constructed independently for three distinct cortical metrics: cortical curvature, surface area and grey matter volume using the same time-based linear mixed-effects model described for the functional analysis.

To justify the choice of the full random-intercept and random-slope specification, a vertex-wise Log-Likelihood Ratio (LR) test compared the full model against the nested random-intercept-only model. The resulting significance map was corrected for multiple comparisons across both hemispheres concurrently using the Benjamini-Hochberg False Discovery Rate (FDR) procedure using a threshold of p< 0.05. Contiguous clusters of surviving vertices were defined on the *fsaverage* surface mesh using *mri_surfcluster* (annotated via the Destrieux atlas, *aparc.a2009s*). A cluster-extent threshold was applied, restricting retention only to cortical clusters with a spatial surface area of at least 50mm2.The surviving surface labels were transformed into 3D volumetric MNI305 space using *fsaverage* (*mri_label2vol*). To facilitate group comparison and standardised reporting, a rigid registration was calculated between the *fsaverage* anatomy and the standard MNI152 template (ICBM MNI152 Non-linear 6th Generation Asymmetric) using *mri_coreg*. The 3D volumetric clusters were then warped into this MNI standard space using the calculated registration data with nearest-neighbour interpolation via *mri_vol2vol*.

To examine whether longitudinal changes in brain anatomy were associated with changes in behavioural performance, anatomical measures from the resulting cortical ROIs were combined with participant-level drift diffusion model estimates of drift rate (v) across sessions in two complementary analyses: First we fitted a linear mixed-effects model separately for each anatomical ROI to test whether session-to-session changes in anatomy were associated with behavioural performance. Anatomical measurements were person-mean centered and session was mean-centered and entered as a continuous predictor. Secondly, we investigated whether baseline anatomical measurements predicted subsequent changes in behavioural performance by fitting ordinary least-squares regression model separately for each ROI with behavioural change as the dependent variable. Behavioural change was defined as the drift rate *v* at the latest available follow-up session minus the baseline drift rate. Analyses were performed separately for each anatomical measure with p-values for the interaction corrected for multiple comparisons across ROIs using the false discovery rate (FDR).

## Data availability

The data collected for this study is available via the research repository Edmond of the Max Planck Society (https://doi.org/10.17617/3.BZW4F0).

## Code availability

All code used for data analysis is available from GitHub (https://github.com/Algebreaker/COCOA).

## Acknowledgements

This work was supported by the German Research Foundation (499339749). The funders had no role in study design, data collection and analysis, decision to publish or preparation of the manuscript.

## Author contributions

Conceptualisation: MAS; Data Acquisition: JF; Formal analysis: JF, GJ; Funding acquisition: MAS; Methodology: JF, GJ, MAS; Software: JF, GJ; Visualisation: JF, GJ; Writing: original manuscript: JF; Writing: review & editing: JF, GJ, MAS

## Ethics declarations

The authors declare no competing interests.

---

**Figure S2.**
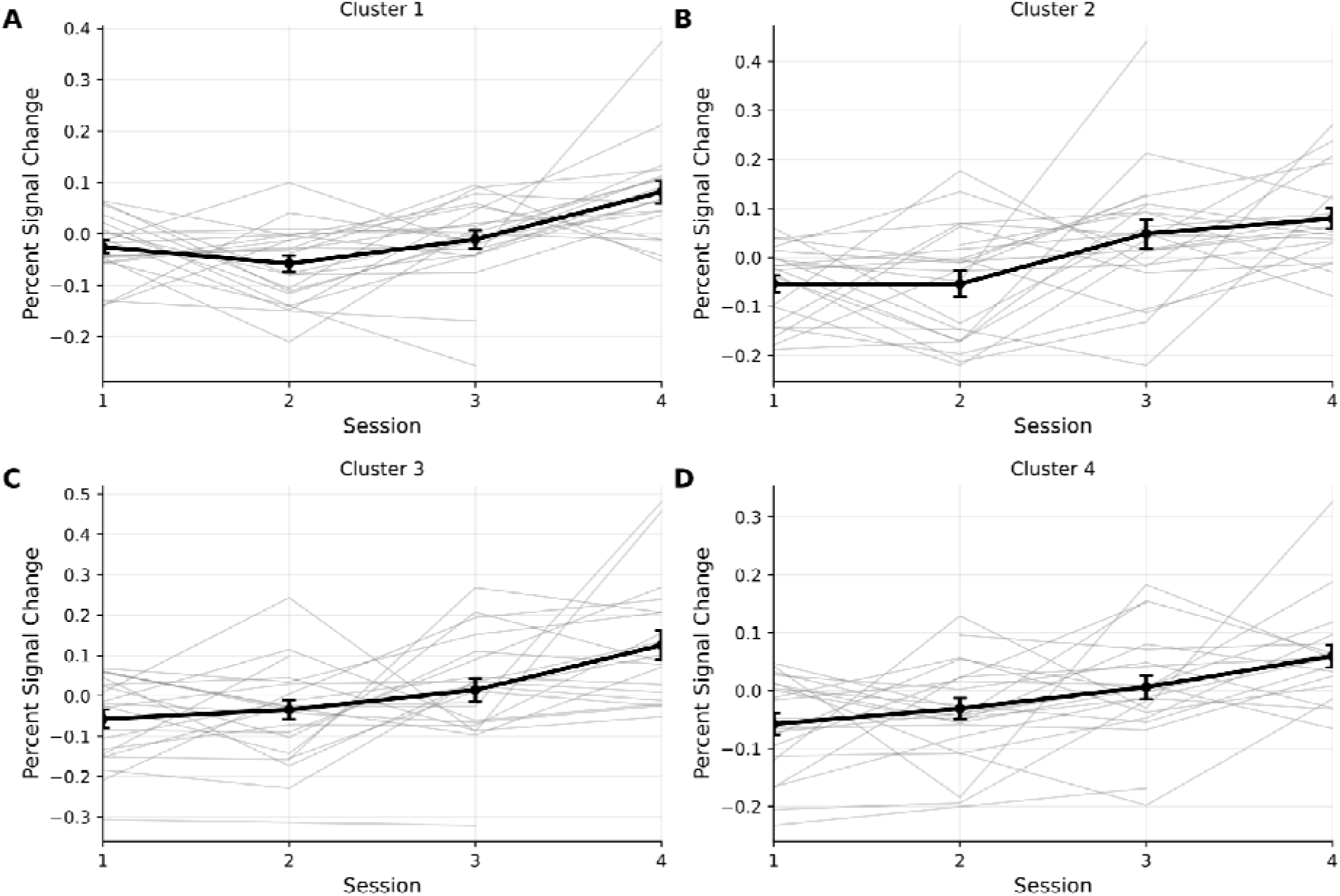
Percent Signal Change in Clusters 1-4. A) Anterior supramarginal gyrus B) Frontal Pole C) Central opercular cortex D) IFG pars triangularis

**Table S1.** Linear mixed-effects model results for within-person associations between functional measures and behavioural performance.

| Metric | CI | $\beta$ within-person anatomy | FDR p |
| --- | --- | --- | --- |
| Anterior supramarginal gyrus | [-2.640, 1.334] | -0.653 | 0.899 |
| Frontal pole | [-1.435, 1.635] | 0.100 | 0.899 |
| Central opercular cortex | [-1.207, 1.606] | 0.199 | 0.899 |
| Inferior frontal gyrus - pars triangularis | [-2.791, 1.556] | -0.618 | 0.899 |

**Table S3.** Linear mixed-effects model results for within-person associations between anatomical measures and behavioural performance.

| Metric | Label | CI | $\beta$ within-person anatomy | FDR p |
| --- | --- | --- | --- | --- |
| Curvature | Middle temporal gyrus (right) | [-41.187, 8.793] | -16.197 | 0.449 |
|  | Inferior temporal gyrus (right) | [-31.853, 5.826] | -13.013 | 0.449 |
|  | Middle occipital and lunate sulci (right) | [-49.092, 17.747] | -15.673 | 0.656 |
|  | Superior occipital and transverse occipital sulci (right) | [-34.037, 64.379] | 15.171 | 0.798 |
|  | Central sulcus (right) | [-17.478, 31.224] | 6.873 | 0.798 |
|  | Parieto-occipital sulcus (right) | [-25.721, 22.775] | -1.473 | 0.905 |
|  | Postcentral gyrus (right) | [-22.464, 4.242] | -9.110 | 0.449 |
|  | Middle temporal gyrus (left) | [-42.059, 0.398] | -20.831 | 0.449 |
|  | Inferior temporal gyrus (left) | [-9.733, 11.836] | 1.052 | 0.905 |
|  | Superior temporal sulcus (left) | [-30.031, 44.139] | 7.054 | 0.867 |
|  | Postcentral sulcus (left) | [-20.252, 3.517] | -8.368 | 0.449 |
| Area | Posterior sylvian fissure (left) | [-0.174, 0.060] | -0.084 | 0.132 |
|  | Superior temporal sulcus (left) | [-0.113, 0.046] | -0.034 | 0.404 |
| Volume | Superior temporal sulcus (left) | [-0.033, 0.015] | -0.009 | 0.478 |

**Table S4.**
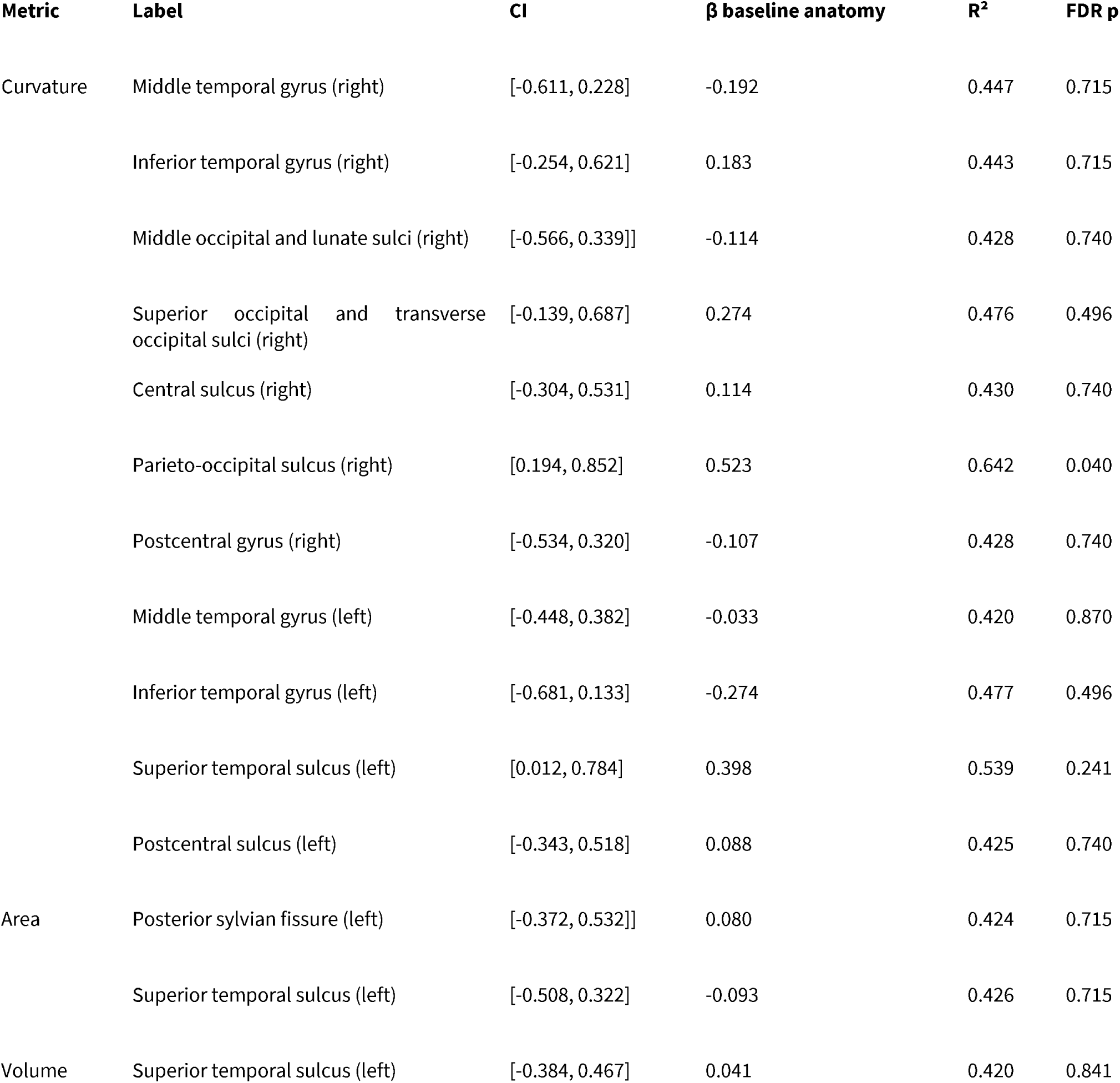
Ordinary least-squares regression results for associations between baseline anatomy and subsequent behavioural change

**Table S5.** Included subjects, sessions and number of valid runs

| Subject | Session | Date & time | Valid runs |
| --- | --- | --- | --- |
| 01 | 1 | 2023-08-14 14:25:58 | 3 |
| 01 | 2 | 2023-10-02 18:00:31 | 4 |
| 01 | 3 | 2024-02-15 18:02:41 | 5 |
| 01 | 4 | 2024-07-23 18:58:36 | 4 |
| 02 | 2 | 2023-10-18 16:17:41 | 2 |
| 02 | 3 | 2024-02-20 18:09:33 | 4 |
| 02 | 4 | 2024-06-21 14:58:42 | 4 |
| 03 | 1 | 2023-08-25 15:17:36 | 4 |
| 03 | 2 | 2023-10-16 16:19:01 | 3 |
| 03 | 3 | 2024-02-26 16:16:10 | 4 |
| 03 | 4 | 2024-06-25 16:24:08 | 4 |
| 04 | 1 | 2023-08-15 17:52:41 | 4 |
| 04 | 2 | 2023-10-09 17:45:48 | 4 |
| 04 | 3 | 2024-02-13 11:51:07 | 4 |
| 04 | 4 | 2024-06-11 16:37:25 | 4 |
| 05 | 1 | 2023-07-14 16:01:28 | 2 |
| 05 | 2 | 2023-10-02 14:45:48 | 4 |
| 05 | 3 | 2024-02-20 16:26:55 | 5 |
| 05 | 4 | 2024-06-20 14:54:57 | 4 |
| 06 | 1 | 2023-07-19 14:12:44 | 4 |
| 06 | 2 | 2023-10-12 10:13:59 | 4 |
| 06 | 3 | 2024-02-23 14:50:28 | 4 |
| 06 | 4 | 2024-06-24 16:00:24 | 4 |
| 07 | 1 | 2023-08-09 14:57:56 | 4 |
| 07 | 2 | 2023-10-02 08:49:54 | 2 |
| 07 | 3 | 2024-02-23 17:22:08 | 2 |
| 08 | 1 | 2023-08-17 15:35:22 | 4 |
| 08 | 2 | 2023-10-02 16:29:28 | 4 |
| 08 | 3 | 2024-02-16 16:25:56 | 4 |
| 08 | 4 | 2024-06-14 17:42:04 | 4 |
| 09 | 1 | 2023-08-07 15:22:32 | 4 |
| 09 | 2 | 2023-10-09 14:50:28 | 2 |
| 09 | 3 | 2024-03-15 16:33:39 | 4 |
| 09 | 4 | 2024-06-20 13:15:31 | 4 |
| 10 | 1 | 2023-08-15 14:57:45 | 2 |
| 10 | 2 | 2023-10-12 12:11:59 | 2 |
| 11 | 1 | 2023-08-18 16:51:55 | 4 |
| 11 | 2 | 2023-10-02 10:09:11 | 3 |
| 11 | 3 | 2024-04-02 16:52:16 | 4 |
| 11 | 4 | 2024-06-12 16:32:35 | 3 |
| 12 | 1 | 2023-08-14 16:04:48 | 4 |
| 12 | 2 | 2023-10-13 16:22:22 | 4 |
| 12 | 3 | 2024-02-28 17:41:37 | 3 |
| 12 | 4 | 2024-06-14 16:23:48 | 4 |
| 13 | 1 | 2023-07-31 15:19:45 | 4 |
| 13 | 2 | 2023-10-06 14:49:05 | 4 |
| 13 | 3 | 2024-02-20 14:50:04 | 4 |
| 13 | 4 | 2024-09-10 16:24:12 | 4 |
| 14 | 1 | 2023-07-19 08:52:53 | 4 |
| 14 | 2 | 2023-10-05 16:09:12 | 4 |
| 14 | 3 | 2024-02-23 10:16:43 | 4 |
| 14 | 4 | 2024-06-13 16:18:21 | 4 |
| 15 | 1 | 2023-07-28 13:53:15 | 2 |
| 15 | 2 | 2023-10-12 13:42:55 | 3 |
| 15 | 4 | 2024-06-21 16:12:59 | 3 |
| 16 | 1 | 2023-08-25 16:57:40 | 3 |
| 16 | 2 | 2023-10-13 17:43:41 | 4 |
| 16 | 3 | 2024-03-13 16:20:39 | 4 |
| 16 | 4 | 2024-07-19 14:29:08 | 5 |
| 17 | 2 | 2023-10-02 13:13:09 | 4 |
| 17 | 3 | 2024-02-21 16:18:30 | 4 |
| 17 | 4 | 2024-06-20 11:52:23 | 4 |
| 18 | 1 | 2023-08-15 16:45:18 | 4 |
| 18 | 2 | 2023-10-13 11:05:58 | 4 |
| 18 | 3 | 2024-02-20 10:38:41 | 4 |
| 18 | 4 | 2024-06-12 18:08:40 | 4 |
| 19 | 1 | 2023-07-17 16:51:58 | 3 |
| 19 | 3 | 2024-02-13 17:30:57 | 4 |
| 19 | 4 | 2024-06-26 08:10:06 | 4 |
| 20 | 1 | 2023-08-16 17:15:23 | 3 |
| 20 | 2 | 2023-10-06 17:42:50 | 4 |
| 20 | 3 | 2024-02-26 17:15:23 | 4 |
| 21 | 1 | 2023-08-21 15:21:58 | 2 |
| 21 | 2 | 2023-11-02 16:38:39 | 4 |
| 21 | 3 | 2024-03-07 18:21:07 | 4 |
| 21 | 4 | 2024-06-10 17:43:16 | 2 |
| 22 | 1 | 2023-07-19 13:07:06 | 2 |
| 22 | 3 | 2024-02-13 15:56:03 | 3 |
| 23 | 1 | 2023-07-28 16:44:53 | 4 |
| 23 | 2 | 2023-10-06 16:18:35 | 4 |
| 23 | 3 | 2024-02-15 16:35:13 | 4 |
| 23 | 4 | 2024-06-20 10:20:21 | 3 |

